# Embryonic environment impacts adult traits and phenotypic integration in the pea aphid

**DOI:** 10.64898/2026.01.14.699474

**Authors:** Lauren E. Gregory, Julia G. McDonough, W. Anthony Frankino, Jennifer A. Brisson

## Abstract

Understanding how the embryonic environment affects adult phenotype is critical in the context of both human health and the increasing variance of environmental conditions due to climate change. Here we consider this topic within the framework of phenotypic integration, i.e., the pattern of correlations among elements of functionally robust trait suites. We investigated the life-long consequences of alternative embryonic environments on phenotypic integration in the pea aphid (*Acyrthosiphon pisum*). We exposed live-bearing pea aphid mothers to an adverse, high-density environment or a benign, low-density environment and measured morphological traits, fecundity, and transcriptional profiles in the resulting winged and wingless adult offspring. We observe that morphological integration decreases in both morphs in response to the adverse maternal environment. Transcriptional integration, on the other hand, shows a morph-specific response: integration increases in wingless offspring but decreases in winged offspring in response to the adverse maternal environment. Our results show the remarkable phenotypic diversity that a single genotype can express in response to environmental variation; that maternal environmental conditions can have strong effects on offspring trait variation and trait integration; and that these effects differ at the morphological and transcriptional levels.

## Introduction

Exposure to environmental variation during development can have long-term, even intergenerational, effects on adult phenotypes [1–3]. This phenomenon, called phenotypic plasticity, is the environment-specific expression of phenotypic variation; such plasticity is often genotype-specific and can generate continuous or discrete trait values [4–6]. Plasticity can be active and adaptive, where individuals respond to environmental cues by expressing environment appropriate, fitness-enhancing phenotypes. For example, water fleas (*Daphnia spp)* respond to predator cues: mothers detecting predator chemical cues signal to their developing offspring, resulting in the expression of fitness-enhancing anti-predator morphologies, behaviors, and life-histories in progeny [7, 8]. Alternatively, plasticity can be passive or nonadaptive, as with prenatal exposure to low nutrition in humans that produces offspring with atypical proportions at birth and higher incidences of systemic health issues well into adulthood [9, 10]. In either case, plasticity plays a crucial role in shaping the evolutionary potential of populations in patchy, changing, or novel environments [11].

Plasticity is often studied on a trait-by-trait basis (e.g., [12]) but organisms are composed of integrated, modular units that often respond to the environment with simultaneous changes across multiple traits. Individual traits may or may not change in concert with other traits. It’s therefore critical to consider plasticity in a multivariate context [13]. If traits are correlated and vary along the same axis, then they may respond to selection together; this can be advantageous if the joint optima is aligned with the axis of covariation or constraining if it is not [14, 15]. If they do not covary, then this is a potential barrier for responding to environmental pressures via directional selection [16].

One way to study correlated phenotypic variation is to consider the degree of morphological integration [17], which describes the pattern, strength, and resiliency of trait correlations. Traits can be correlated due to reasons such as shared developmental mechanisms or biomechanical constraints, among others. Strong integration can result in robust sets of traits that are buffered against external perturbations, although extreme stress can result in the breakdown of that buffering, the release of cryptic genetic variation, and the increase of phenotypic variation [18–20]. The degree of morphological integration, therefore, is itself context-dependent, meaning that integration may be stronger in one environment compared to another. Moreover, a greater degree of morphological integration can result in higher fitness for an organism, if integration is related to function [21].

Our focus here is on the long-term consequences of benign and adverse maternal environments on offspring phenotypic integration in the pea aphid, *Acyrthosiphon pisum*. Pea aphids have long been studied for their transgenerational wing plasticity, in which adult asexual females produce genetically identical winged or wingless daughters when the mothers experience high-quality or low-quality conditions, respectively [22]. Low-quality maternal conditions largely include high population densities, although they can also include poor nutrition and/or the presence of predators [22–24]. This plasticity occurs during the spring and summer months, when aphids are viviparous and asexual, giving live birth to genetically identical daughters via a modified meiosis that bypasses recombination [25]. Aphid mothers process their environments and signal to the embryos in their ovaries, in a process likely mediated by biogenic amines and hormones [26–28].

The pea aphid plasticity, as with other adaptive plasticities, results in environment-phenotype matching: wingless aphids in benign conditions maximize reproduction, while winged aphids can disperse away from an adverse environment. This is a reproductive-dispersal trade-off commonly observed in wing dimorphic insects [29]. Differences between the winged and wingless morphs are systemic, with winged morphs having more complex sensory systems that aid them in finding more suitable hosts on which to settle [30–32], while wingless morphs have an increased rate of development [33, 34], and both morphs differ extensively in their transcriptional profiles [35–38]. Smaller-scale trait size differences also exist, with wingless generally larger than winged morphs [39]. Yet despite the broad environment-morph matching that occurs, most pea aphid lines produce both winged and wingless morphs in both environments; it is the proportion of each morph that changes in response to benign compared to adverse conditions [40, 41].

The pea aphid wing plasticity therefore offers a unique opportunity to investigate trait integration in response to benign compared to adverse environments: pea aphids produce both morphs in each environment, morphs are genetically identical, and morphs comprise evolved, systemic differences. Here we exposed pea aphid mothers to a benign (low density) or adverse (high density) environment and then reared offspring to adulthood in a common, benign environment. At adulthood, we assessed morphological, reproductive, and gene expression traits in both winged and wingless morphs. Importantly, we used a single genotype (*i.e.,* genetically identical individuals) for the entirety of the study, which allowed us to attribute all trait variation to environmental effects. Within each morph, we compared trait integration between the two maternal conditions, hypothesizing that adverse conditions would decrease trait integration at both transcriptional and morphological levels.

## Materials and Methods

### Overview

Our overall experimental protocol was to raise aphids to adulthood, apply an experimental treatment to the adult asexual females for 24hrs, grow their progeny in a common environment, measure fitness traits at reproductive maturity, and then process adult carcasses for morphological measures or RNA sequencing. Because asexual females are viviparous, the aphid mothers and their soon to be born embryos simultaneously experienced the treatment.

### Maternal treatments

We maintained asexual females from a single genotype (line 584) in cages containing *Vicia faba* seedlings at 18°C on a 16:8 (light: dark) cycle. Aphids were reared at low density (3-4 aphids/cage) to reproductive maturity for at least three generations and then wingless adults were randomly assigned to either low- (LD; remained on original low-density plants) or high-density (HD; 12 adults crowded in a 30mm x 15mm diameter Petri dish) conditions for 24hr. Adult aphids were then placed onto leaf plates (a single *Vicia faba* leaf embedded in agar on a 100mm x 15mm Petri dish) for an additional 24hr to reproduce.

### Offspring developmental timing, reproductive, and morphological trait measurements

All nymphs born were moved to individual leaf plates and tracked daily. Aphids were scored upon adulthood as winged or wingless morphs and checked daily for onset of reproductive maturity. The number of nymphs produced by an individual was recorded daily for the first four days of adulthood so that a measure of fecundity could be assessed prior to fixation for morphological measures. After day four of adulthood, we measured the wet mass of each adult, dissected embryos, and counted the number of ovarioles and mature embryos (embryos with pigmented eyes) for each aphid.

Following dissection of embryos from adult offspring (above), we placed the adult carcasses individually into 1.7mL Eppendorf tubes containing 0.7 mL Oudeman’s fluid (5 parts glycerin, 87 parts 70% EtOH, 8 parts glacial acetic acid) for one month. After treatment in 9.1% KOH at 38° C for 20 minutes, we gently scraped out the internal contents of the aphid in a glass well plate. Cleared aphid samples were then moved to a 50% EtOH solution at room temperature for 30m, followed by 70% EtOH for 30m, and 95% EtOH for 1-4h. Aphids were dried by carefully placing them on a KimWipe to allow moisture absorption. Dried, cleared aphids were then placed in clove oil for a minimum of 2h. Samples were mounted in two drops of 50% Canada balsam:50% orange oil and topped with a glass cover slip and sealed with clear nail polish.

We used Fiji to measure metrical morphological traits [42].The calibration was set using a photograph of a 1mm calibration slide taken at each imaging session with a 2.5x objective, so the measured pixel values of the images could be transformed into mm. We used the segmented line tool to measure the lengths of the following linear traits: rostrum length, head width, length of antennal segment 3, length of antennal segment 4, length of front femur, length of front tibia, length of middle femur, length of middle tibia, length of hind femur, length of hind tibia, and cornicle lengths. A meristic trait, number of rhinaria, was measured by imaging slides with a 10x objective. Meristic and metrical traits were measured on three separate occasions by two users (two replicates by one user and one replicate by the other) and the average of repeated measures was used in subsequent analyses. We compared morphological (metrical and meristic listed above) and reproductive (number of ovarioles, mature embryo counts, and reproductive output over the first two days of reproductive maturity) traits within and across morphs using a two-way ANOVA with Type III Sum of Squares, using the ‘anova’ function in the R package car (v3.1.3; [43]). Following significant ANOVA results (p<0.05), pairwise comparisons between group means were performed using the R package emmeans (v.1.11.0; [44]). p-values were adjusted for multiple comparisons using the Tukey method.

Metrical data was standardized to account for body size differences. We found that front femur length scaled linearly with wet mass for each of our four treatment-morph groups (Fig. S1) and therefore used front femur length as a proxy for body size. We performed linear regression on front femur length and every other morphological trait, individually. The slope of each linear regression was used for size standardization using a modified allometric equation (equation 4 of [45]: Y_std_ = Y_obs_(M_avg_/M_obs_)*^b^*. This equation calculates the size standardized measurement (Y_std_), using the recorded, non-size adjusted trait measurement (Y_obs_), the average body size across all specimens (average front femur length, M_avg_), the body size of the individual (front femur length here, M_obs_), and the slope for the linear regression of that trait against body size (*b*). As reproductive data differ in unit of measure, this size standardization was used only for metrical morphological measures.

### Analyses of Phenotypic Integration

Correlation matrices (Kendall’s tau) and corresponding p-value matrices were created and visualized for each maternal treatment-phenotype group (LD winged, HD winged, LD wingless, and HD wingless) using R v3.17 [46], the ggcorrplot package (v0.1.4.1; [47]), and the evolqg package (v0.3.4; [48]), with i) size-standardized morphological measures and reproductive data, or ii) gene expression values as input. Eigenvalue variance was calculated from each correlation matrix and eigenvalue variance distributions for each treatment-morph group were obtained from bootstrap resampling (iterations=1000 for morphological matrices; 100 for transcript matrices) using the R package boot (v1.3.31; [49, 50]). Group eigenvalue variance were compared using a one-way ANOVA, followed by Tukey’s honestly significant difference (HSD) test using the R package stats (v4.3.1; [46]). Variances were compared using a Levene’s test [51].

### RNA-Seq Experimental Design

The same maternal treatments (LD or HD conditions) as above were used, and treated mothers were allowed to reproduce for 24hr post-treatment and were then discarded. Nymphs produced during the 24hr reproductive period were placed on individual leaf plates and monitored daily for molting. Fecundity was tracked over the first two days of reproductive maturity. Ovaries were dissected on day three of reproductive maturity, and number of ovarioles and mature embryos were recorded. Immediately after dissection, carcasses were individually flash frozen in liquid nitrogen, pulverized with a pestle, and stored in TRIzol reagent (Invitrogen) at -80°C. Individual aphid RNA was extracted using a standard TRIzol-chloroform extraction followed by a DNase I treatment (Zymo) and the Zymo RNA Clean & Concentrator-5 kit following the manufacturer’s protocol.

### RNA Sequencing and Analysis

cDNA library preparation and RNA sequencing were conducted by Genewiz (South Plainfield, NJ, USA) using Illumina HiSeq 150 bp paired end reads. Raw reads are in the processing stage post-submission to SRA (BioProject/BioSample accession not yet assigned). Raw reads were trimmed using Trimmomatic v0.39 and quality was checked post-trimming using MultiQC v1.14. Read mapping was done using HISAT2 v2.2.1 and the v3.0 pea aphid genome. samtools and bowtie2 (versions 1.17 and 2.5.1, respectively) were used to sort and index mapped reads. HTSeq-count v2.0.3 was used to count reads. We performed differential expression analyses using DESeq2 with variance-stabilizing transformation (vst), incorporating bash scripts and R Studio v3.17 for these analyses. P-values were corrected using the Benjamini-Hochberg method. We performed GO enrichment analysis using g:GOSt of g:Profiler [52] with background set to the pea aphid genome GCF_005508785.1 (v2), statistical domain scope set to ‘only annotated genes’, significance threshold set to Bonferroni correction, and user threshold set at ‘0.05’. Weighted gene co-expression network analysis was conducted using the R package WGCNA (v1.73; [53, 54]). We excluded any genes that had less than 15 counts in fewer than 75% of samples. Regularized log transformed counts were used. Gene co-expression matrices were built using Pearson correlations. The network was constructed using a soft power threshold of 16, which agreed with a scale-free topology (Fig. S2). Modules were determined in a single block. We used the ‘dynamicTreeCut’ function with deepSplit set to 2, mergeCutHeight set to 0.25, and a minimum cluster size set to 30 to merge similar modules. A signed adjacency matrix was computed and transformed to a topological overlap measure (TOM) matrix and trait relationships per module were determined. The corPvalueStudent function was used to calculate the Student asymptotic p-value.

## Results

### Maternal environment impacts offspring adult traits

We found that maternal environment (HD or LD) impacted adult offspring reproductive and morphological characters in a morph-specific manner (Fig. 1A-C). Most variation in these traits predictably occurred between the winged and wingless morphs (blue versus green clouds, Fig. 1A), however, examination of the morphs separately revealed systemic differences based on maternal treatment in wingless (Fig. 1C) but not winged (Fig. 1B) individuals. In particular, the phenotypes of wingless offspring from high density (HD) mothers shifted towards those of the winged morph (Fig.1A).

**Figure 1.**
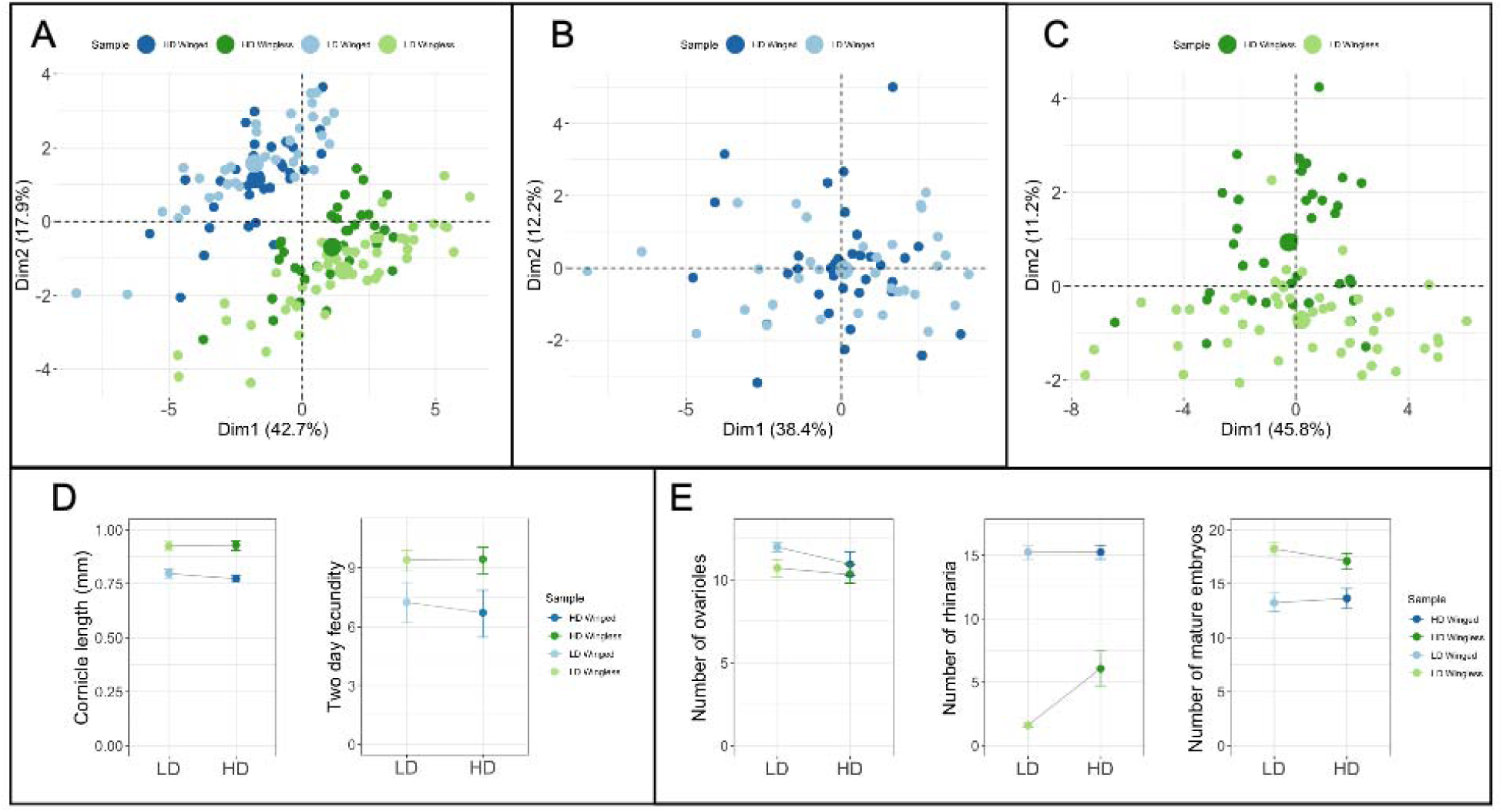
Maternal environment impacts reproductive and morphological characters in a morph-specific manner. A) Principal component analysis (PCA) of morphological and reproductive measures. Morphological data include rostrum length, head width, length of antennal segments 3&4, length of each femur and tibia, cornicle length, and number of rhinaria. Reproductive data include number of mature embryos and number of ovarioles. Centroids for each group are shown as enlarged circles of the respective group color-coding. B) PCA of morphological and reproductive measures for winged and C) wingless samples only. D) Reaction norms for traits that differ significantly between morphs, but not between treatments.

We also examined trait reaction norms on a trait-by-trait basis. We discovered that several traits differed between the morphs (Tables 1 and 2). For example, the length of cornicles, the abdominal structures that exude alarm pheromone [55], are longer in wingless morphs, but not responsive to maternal treatment (Fig.1D, left). Similarly, fecundity over the first two days of reproductive maturity is higher in the wingless compared to winged morph but is not statistically differentially affected by treatment (Fig. 1D, right).

**Table 1:**
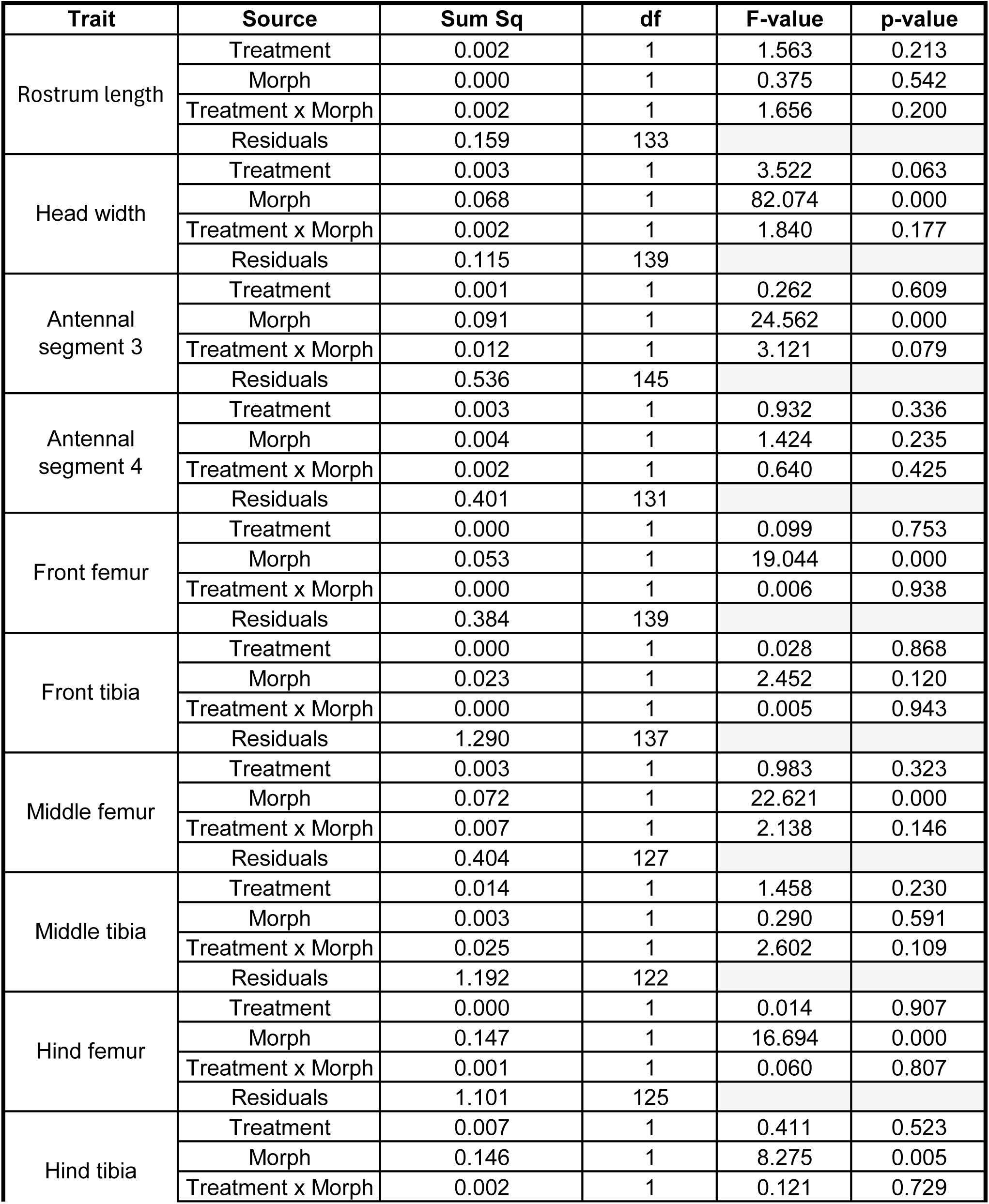
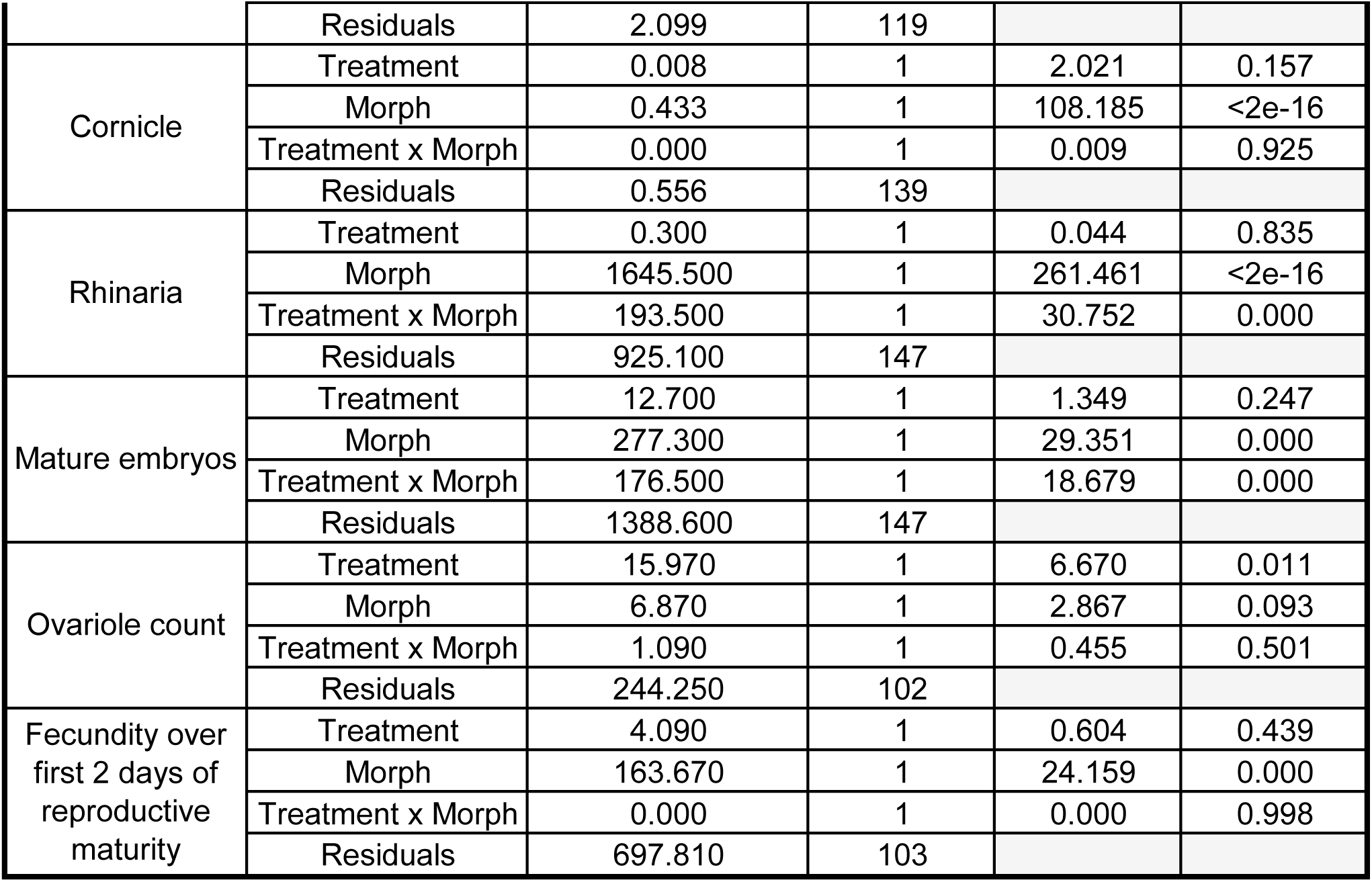
Summary results of two-way ANOVAs testing the effect of maternal treatment (HD or LD) and morph (W or WL) on 12 morphological and reproductive traits. Sum of squares, degrees of freedom (df), F-statistics, and p-values are shown. Analyses were performed using Type III Sum of Squares.

| Trait | Source | Sum Sq | df | F-value | p-value |
| --- | --- | --- | --- | --- | --- |
| Rostrum length | Treatment | 0.002 | 1 | 1.563 | 0.213 |
|  | Morph | 0.000 | 1 | 0.375 | 0.542 |
|  | Treatment x Morph | 0.002 | 1 | 1.656 | 0.200 |
|  | Residuals | 0.159 | 133 |  |  |
| Head width | Treatment | 0.003 | 1 | 3.522 | 0.063 |
|  | Morph | 0.068 | 1 | 82.074 | 0.000 |
|  | Treatment x Morph | 0.002 | 1 | 1.840 | 0.177 |
|  | Residuals | 0.115 | 139 |  |  |
| Antennal segment 3 | Treatment | 0.001 | 1 | 0.262 | 0.609 |
|  | Morph | 0.091 | 1 | 24.562 | 0.000 |
|  | Treatment x Morph | 0.012 | 1 | 3.121 | 0.079 |
|  | Residuals | 0.536 | 145 |  |  |
| Antennal segment 4 | Treatment | 0.003 | 1 | 0.932 | 0.336 |
|  | Morph | 0.004 | 1 | 1.424 | 0.235 |
|  | Treatment x Morph | 0.002 | 1 | 0.640 | 0.425 |
|  | Residuals | 0.401 | 131 |  |  |
| Front femur | Treatment | 0.000 | 1 | 0.099 | 0.753 |
|  | Morph | 0.053 | 1 | 19.044 | 0.000 |
|  | Treatment x Morph | 0.000 | 1 | 0.006 | 0.938 |
|  | Residuals | 0.384 | 139 |  |  |
| Front tibia | Treatment | 0.000 | 1 | 0.028 | 0.868 |
|  | Morph | 0.023 | 1 | 2.452 | 0.120 |
|  | Treatment x Morph | 0.000 | 1 | 0.005 | 0.943 |
|  | Residuals | 1.290 | 137 |  |  |
| Middle femur | Treatment | 0.003 | 1 | 0.983 | 0.323 |
|  | Morph | 0.072 | 1 | 22.621 | 0.000 |
|  | Treatment x Morph | 0.007 | 1 | 2.138 | 0.146 |
|  | Residuals | 0.404 | 127 |  |  |
| Middle tibia | Treatment | 0.014 | 1 | 1.458 | 0.230 |
|  | Morph | 0.003 | 1 | 0.290 | 0.591 |
|  | Treatment x Morph | 0.025 | 1 | 2.602 | 0.109 |
|  | Residuals | 1.192 | 122 |  |  |
| Hind femur | Treatment | 0.000 | 1 | 0.014 | 0.907 |
|  | Morph | 0.147 | 1 | 16.694 | 0.000 |
|  | Treatment x Morph | 0.001 | 1 | 0.060 | 0.807 |
|  | Residuals | 1.101 | 125 |  |  |
| Hind tibia | Treatment | 0.007 | 1 | 0.411 | 0.523 |
|  | Morph | 0.146 | 1 | 8.275 | 0.005 |
|  | Treatment x Morph | 0.002 | 1 | 0.121 | 0.729 |
|  | Residuals | 2.099 | 119 |  |  |
| Cornicle | Treatment | 0.008 | 1 | 2.021 | 0.157 |
|  | Morph | 0.433 | 1 | 108.185 | <2e-16 |
|  | Treatment x Morph | 0.000 | 1 | 0.009 | 0.925 |
|  | Residuals | 0.556 | 139 |  |  |
| Rhinaria | Treatment | 0.300 | 1 | 0.044 | 0.835 |
|  | Morph | 1645.500 | 1 | 261.461 | <2e-16 |
|  | Treatment x Morph | 193.500 | 1 | 30.752 | 0.000 |
|  | Residuals | 925.100 | 147 |  |  |
| Mature embryos | Treatment | 12.700 | 1 | 1.349 | 0.247 |
|  | Morph | 277.300 | 1 | 29.351 | 0.000 |
|  | Treatment x Morph | 176.500 | 1 | 18.679 | 0.000 |
|  | Residuals | 1388.600 | 147 |  |  |
| Ovariole count | Treatment | 15.970 | 1 | 6.670 | 0.011 |
|  | Morph | 6.870 | 1 | 2.867 | 0.093 |
|  | Treatment x Morph | 1.090 | 1 | 0.455 | 0.501 |
|  | Residuals | 244.250 | 102 |  |  |
| Fecundity over first 2 days of reproductive maturity | Treatment | 4.090 | 1 | 0.604 | 0.439 |
|  | Morph | 163.670 | 1 | 24.159 | 0.000 |
|  | Treatment x Morph | 0.000 | 1 | 0.000 | 0.998 |
|  | Residuals | 697.810 | 103 |  |  |

**Table 2:**
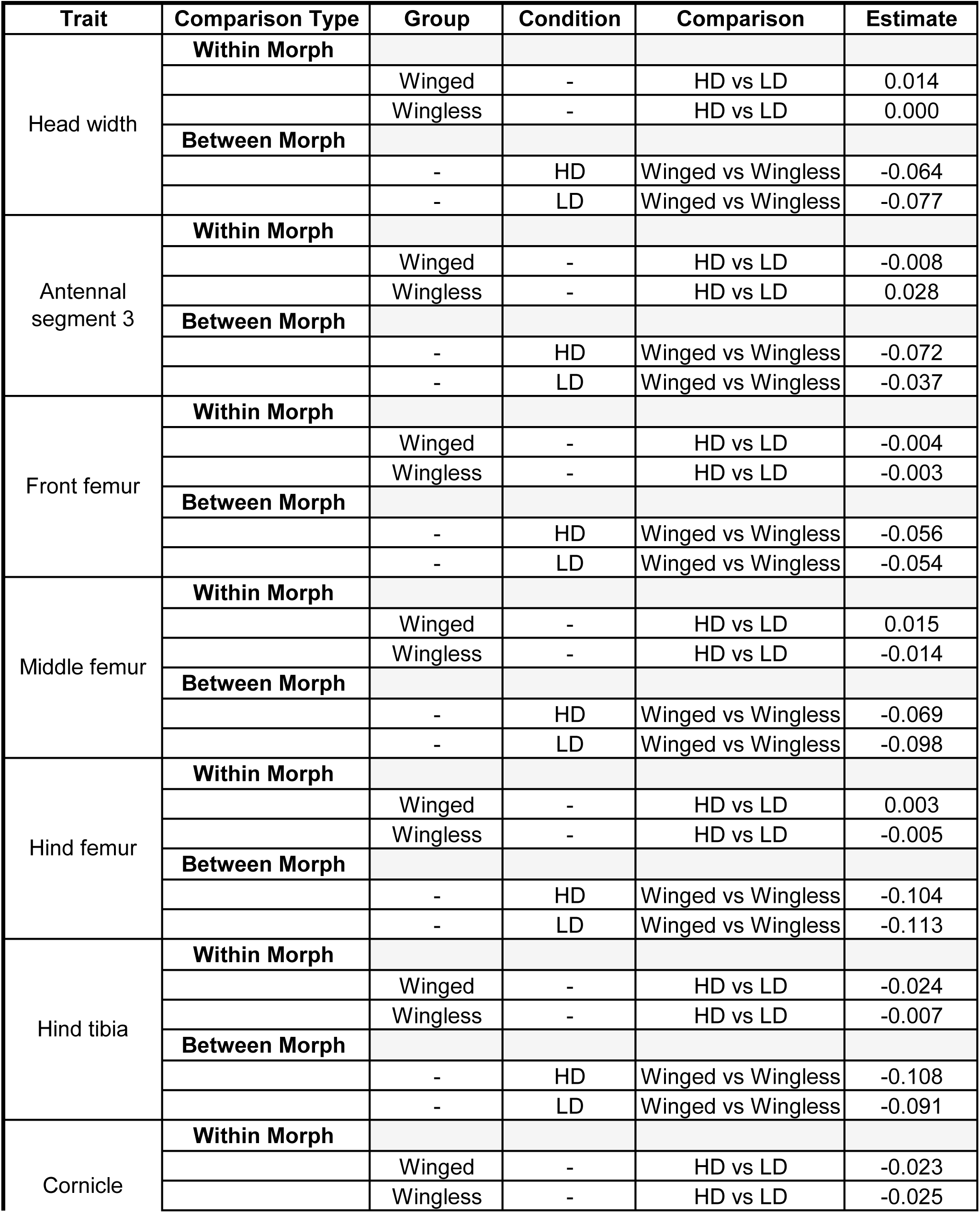
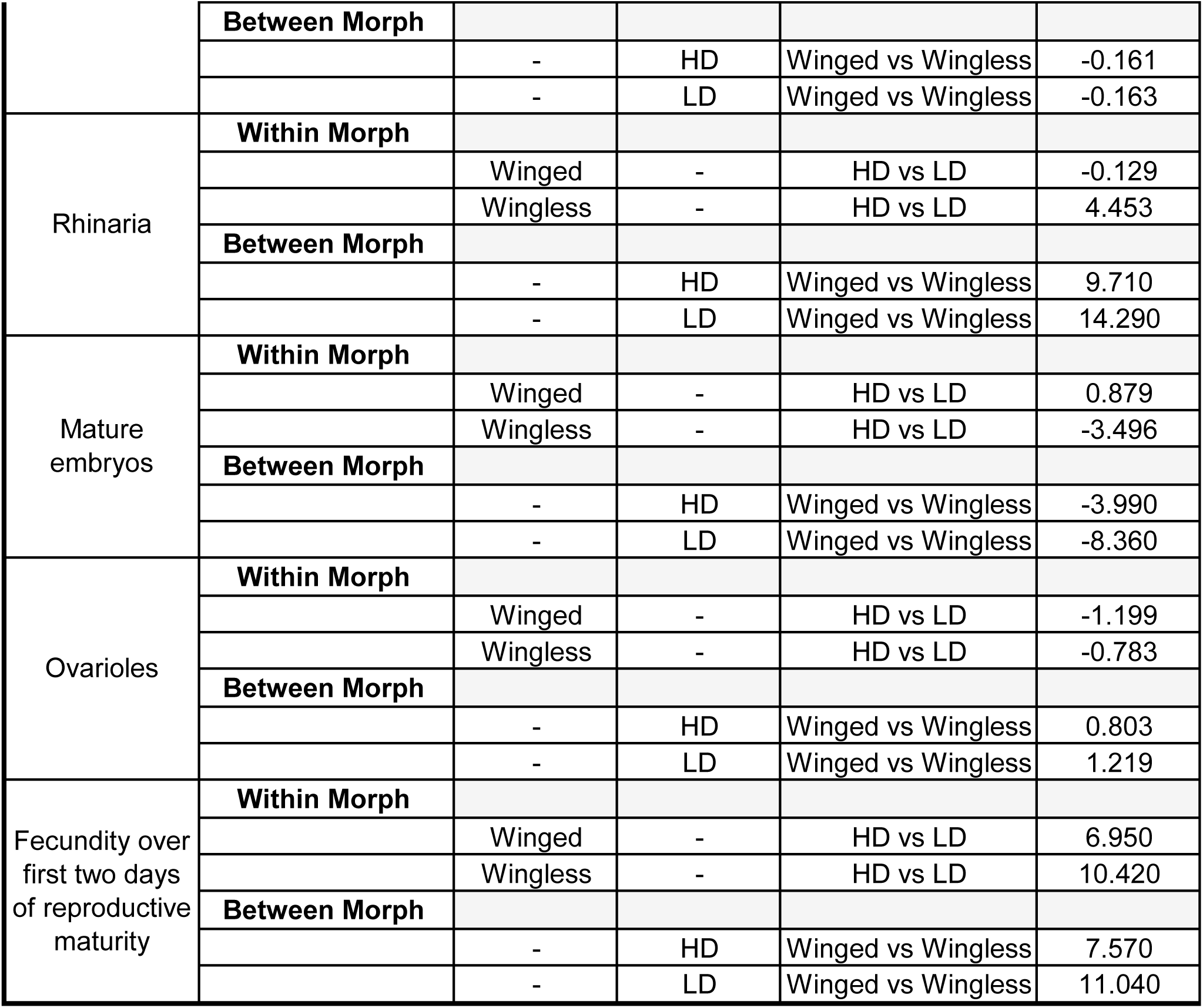
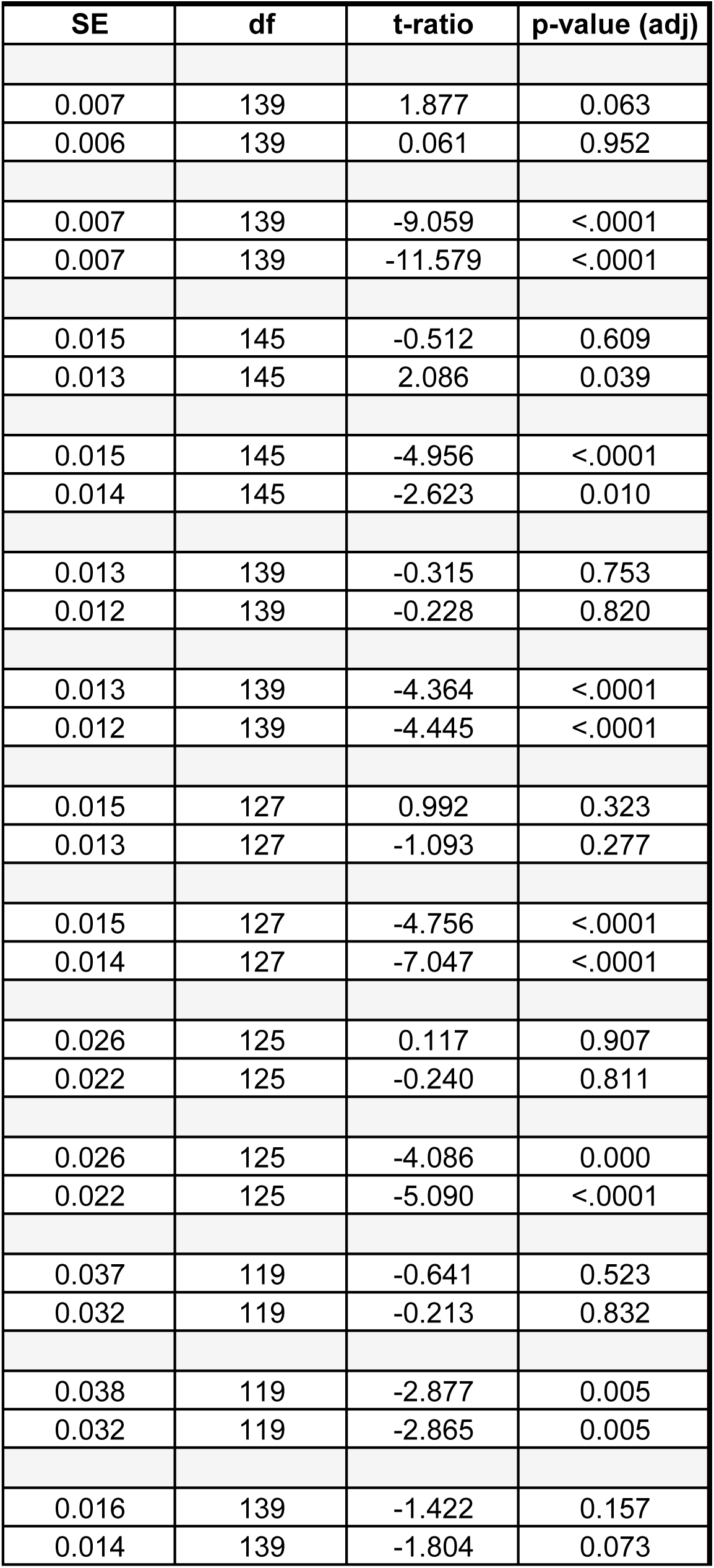
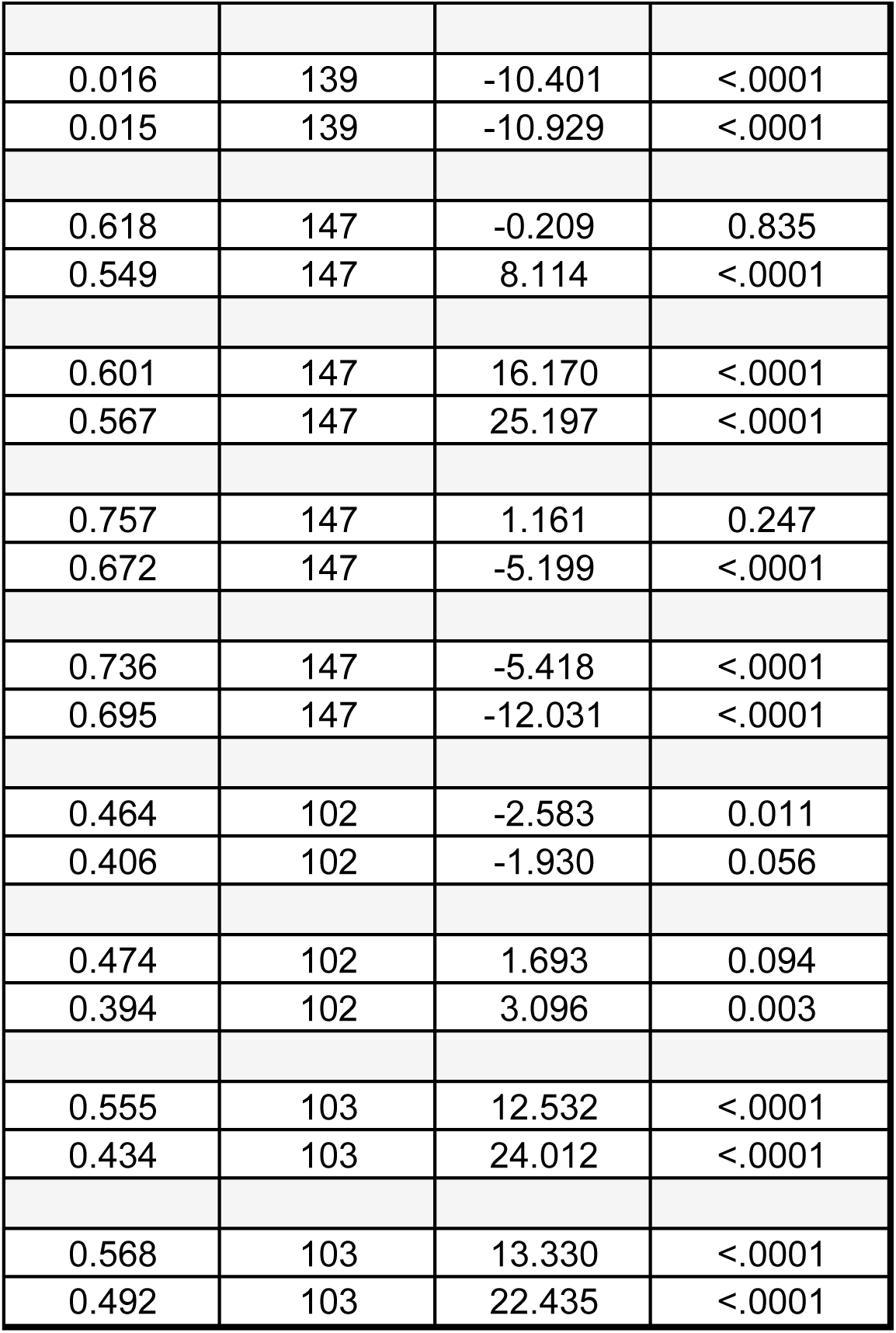
Post-hoc pairwise comparisons of estimated marginal means (EMMs) for morphological and reproductive traits with significant ANOVA effects. Estimates (differences), standard error (SE), degrees of freedom (df), t-ratios, and adjusted p-values are shown. P-values were adjusted for multiple comparisons using the Tukey method. HD = high-density maternal treatment; LD = low-density maternal treatment.

Left plot shows cornicle lengths, while right plot shows the fecundity over the first two days of reproductive maturity. Standard error bars are shown for each sample group. E) Reaction norms for traits that differ significantly between treatment in at least one morph. Left plot shows number of ovarioles, middle plot shows number of rhinaria, and right plot shows number of mature embryos. Standard error bars are shown for each sample group. For all plots, LD = maternal low density and HD = maternal high density. Winged samples are shown in shades of blue and wingless samples are in green. Darker shades for each morph indicate HD maternal treatments.

We also identified traits that responded differently to maternal treatment (Tables 1 and 2). Winged offspring from HD mothers had more ovarioles than winged offspring from LD mothers (Fig. 1E, left). Wingless daughters of HD mothers had more rhinaria – sensory organs on aphid antennae that aid in detecting suitable host plants during dispersal – than those of LD mothers (Fig. 1E, center; [56]). And the number of mature embryos was higher in wingless females from LD compared to HD mothers (Fig. 1E, right).

Overall, these findings demonstrate that the embryonic environment impacts adult trait expression. Plasticity varies among traits, with the wingless morph generally exhibiting a stronger response to maternal density cues than the winged morph.

### Maternal environment impacts offspring phenotypic integration

We next investigated whether maternal environment affected the trait integration across reproductive and morphological characters using two approaches. First, we calculated the eigenvalue variance of the traits; this provides a single value for integration across all traits, with larger values indicating higher levels of integration [57]. Second, we examined pairwise correlations for all traits to highlight trait-by-trait differences.

Preliminary analyses showed that nearly all traits were affected by body size, indicating the importance of this variable on trait integration (Fig. S3). To investigate levels of integration that remained after the effects of size were removed, we size-standardized the morphological data using front femur length as a proxy for body size (see Methods). We found that integration values for size-corrected offspring traits, as measured by eigenvalue variance, were impacted by maternal environment. For both morphs, offspring from LD mothers exhibited significantly higher phenotypic integration than did offspring from HD progeny (Fig. 2A). Moreover, the standard deviation of eigenvalue variance was higher in the morphs with higher integration values, indicating that although they have higher mean integration, they are also more variable in their integration values (Levene’s test [51]; WL morphs F-value = 562.4, p <2.2e-16; W morphs F-value = 489.2, p <2.2e-16).

**Figure 2.**
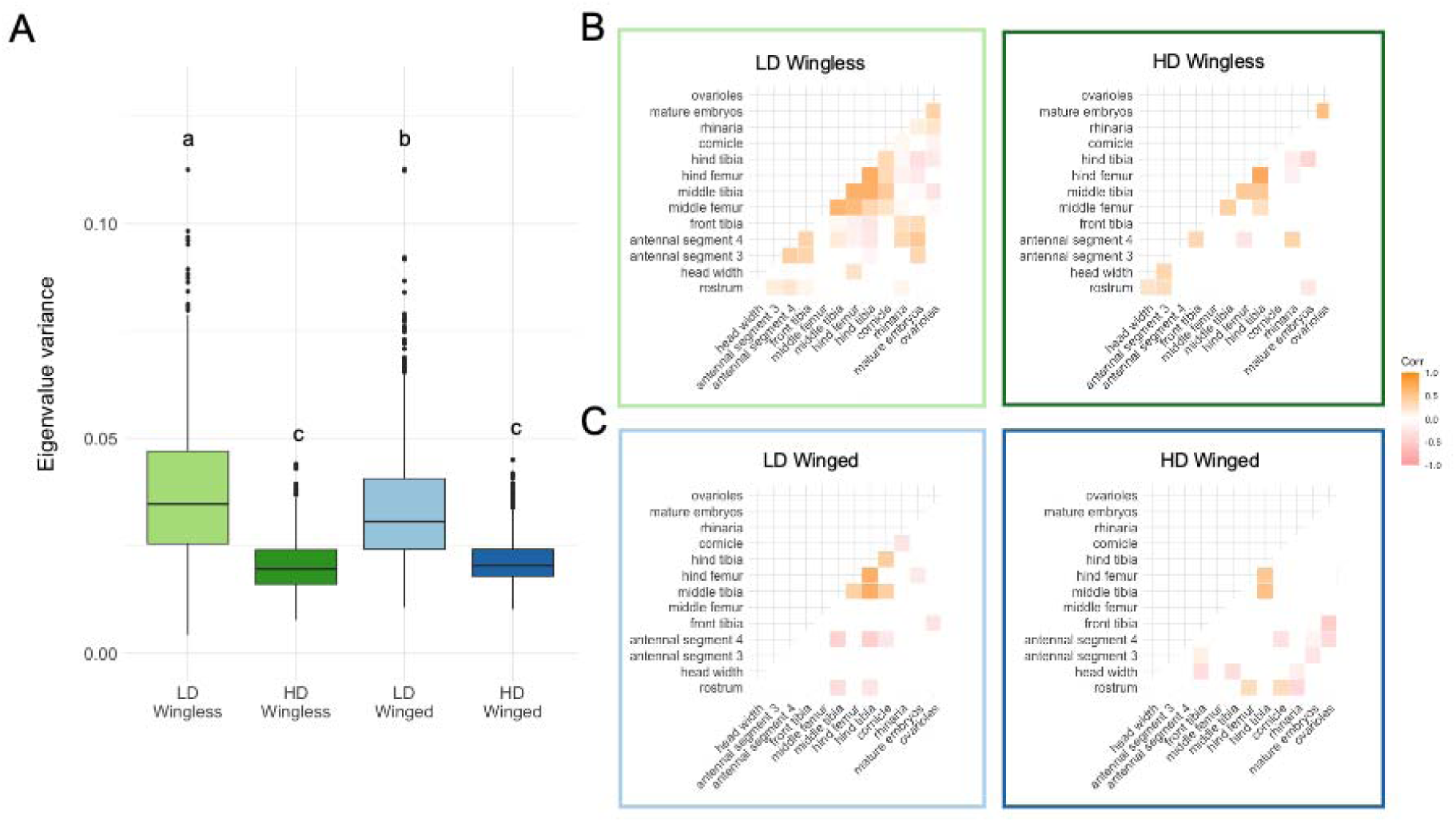
High maternal density reduces offspring morphological and reproductive integration. A) Integration values for morphological and reproductive data shown as eigenvalue variance boxplots. Morphological data are size standardized from front femur lengths. Integration values were calculated using 1000 bootstrap resamples. Results from a one-way ANOVA using treatment-morph as a grouping variable yielded p < 2.2e-16, with post-hoc (Tukey HSD) differences shown. B) Correlograms for wingless samples from LD and HD maternal conditions using Kendall’s tau correlation coefficients. Morphological data are size standardized as in panel A. C) Correlograms for winged samples from LD and HD maternal conditions using Kendall’s tau correlation coefficients. Morphological data are size-standardized and reproductive data are the same as Figure 1A. Darker orange hues indicate a strong positive correlation, while darker pink hues indicate stronger negative correlations. Only significant correlations are shown; non-significant correlations are displayed in white.

Trait-by-trait correlations showed a parallel pattern. In wingless offspring from LD mothers, most size-corrected traits exhibited significant correlations with one another (53 significant of 91 total correlations, Fig. 2B, left). In contrast, wingless offspring from HD mothers had lower numbers of significantly correlated traits (22 significant out of 91 total correlations) and a different pattern of correlations, with some disappearing entirely (Fig. 2B, right). Winged progeny had fewer significant trait correlations compared to wingless progeny, and the strength and number of significant correlations modestly decreased in offspring from HD mothers (18 significant in LD, down to 15 in HD of 91 total correlations, Fig. 2C).

### Morph-specific gene expression profiles differ in response to maternal density environment

We next sought to determine if the morphological and reproductive responses to maternal environment in progeny were accompanied by physiological changes manifested in the progeny’s transcriptomes. As before, we exposed wingless aphid mothers of a single genotype to HD or LD environment for 24h. When progeny reached the third day of adulthood, we removed their ovaries and embryos and collected total RNA from individual carcasses. Our rationale for ovary and embryo removal was that we wanted to observe maternal effects on the progeny generation, not the next generation which would be captured in the embryos.

We found that maternal environment had lasting effects on the physiology of adult progeny. Although we observed the largest transcriptome differences between winged and wingless morphs (Fig. 3A), many genes had plastic expression in response to maternal treatments within morphs (Fig. 3C). In wingless progeny, 122 genes had significantly different transcript levels (adjP < 0.01; Fig. 3C, left; Table S1) between HD and LD maternal treatments, with the majority at higher levels in progeny of HD mothers. Their molecular function gene ontology terms were overrepresented for transmembrane transporter activity. Winged progeny had fewer differences (94 genes, Fig. 3C, right; Table S2), but exhibited the opposite pattern: most transcripts had higher levels in progeny of LD mothers. This gene list had overrepresentation of structural constituent of ribosome, ubiquitin protein-related activity, and aminoacyltransferase activity for molecular function GO terms. Specifically, ribosomal proteins were at higher levels and ubiquitinization-related proteins were at lower levels in HD mothers, indicating an increased protein need in winged progeny from stressful maternal environments, or a decreased protein need in winged offspring from benign maternal environments.

**Figure 3.**
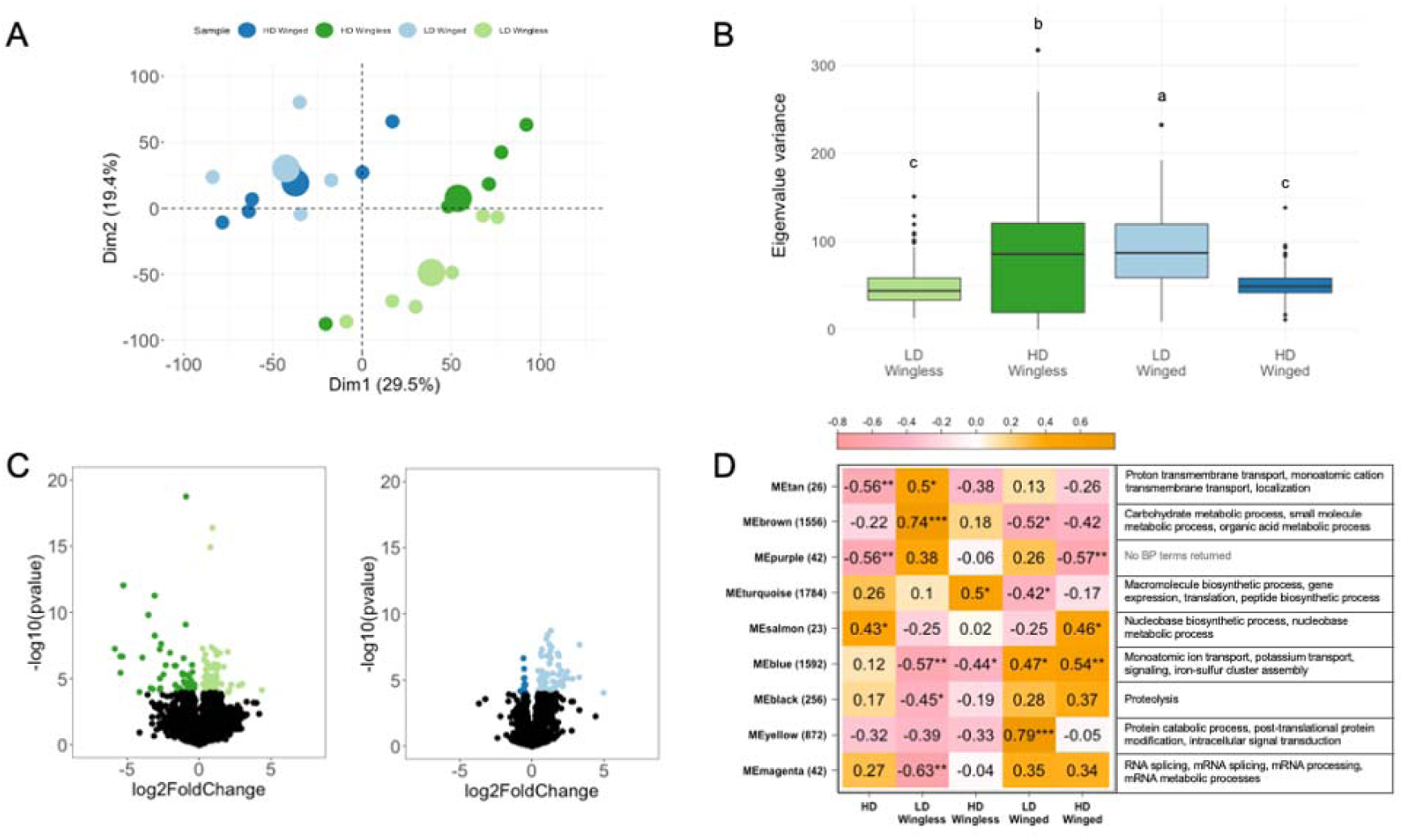
Maternal environment has lasting effects on the physiology of adult progeny. A) PCA of transcript levels with HD winged samples (n=6) shown in dark blue, LD winged samples (n=5) in light blue, HD wingless samples (n=5) in dark green, and LD wingless samples (n=6) in light green. Centroids for each group are shown as enlarged versions of their respective color. B) Transcriptional integration across samples. Integration values were calculated using 100 bootstrap resamples of all transcripts. Results from a one-way ANOVA using treatment-morph as a grouping variable yielded p < 2.2e-16 with post-hoc (Tukey HSD) differences shown. C) Volcano plots for wingless samples (left plot; 11,865 total genes after filtering) and winged samples (right plot; 11,972 total genes after filtering). Colored points indicate that a gene is differentially expressed with an adjusted p-value < 0.01 and is at a higher level in the samples of the same color. Black points are genes that are not significantly differentially expressed. D) Reduced module-trait relationship heatmap from weighted gene co-expression network analysis. Modules are named by color, with the number of genes assigned to each module shown in parentheses. ‘Trait’ groups include all samples from HD maternal groups and each combination of maternal treatment and offspring wing morph. The LD maternal treatment not shown, as it is the binary complement of the HD condition. Orange hues indicate a positive correlation between treatment group or treatment-morph group and module, while pinker hues indicate a negative correlation. Correlation coefficients are shown for each module-trait comparison with the corrected p-value indicated using asterisks. P-values and their asterisks are as follows: p < 0.05 *, p < 0.01 **, p < 0.001 ***. If no asterisk is shown, there is no significant module-trait association. Top gene ontology terms for ‘biological process’ (BP) are listed to the right of the heatmap for each module. Full module-trait heatmap is available in Fig. S4.

Weighted gene co-expression network analysis (WGCNA; [53] showed correlated sets of genes that differed among maternal treatments (tan, brown, turquoise, black and magenta that differed for wingless; brown, purple, turquoise, salmon, and yellow for winged; Fig. 3D). GO enrichment of the significantly correlated gene sets revealed a range of different biological processes (Fig. 3D). The brown module had a striking pattern of 1,556 genes being highly positively correlated in wingless offspring from LD but not HD mothers, with the opposite being true in winged: the 1,556 were highly *negatively* correlated in winged offspring from LD but not HD mothers. This module was enriched for metabolic processes.

We also found that maternal environment affected transcriptional integration in progeny as measured by eigenvalue variance. The pattern differed from that in the combined morphological and reproductive data: rather than maternal crowding causing a decrease in integration in both morphs, we found that maternal crowding decreased integration in winged but not wingless progeny (Fig. 3B). As was the case with morphological integration, the standard deviation of transcriptional eigenvalue variance was higher in the groups with higher integration values (here, HD Wingless and LD Winged, Fig. 3B), (Levene’s test; wingless morphs F-value = 342.3, p <2.2e-16; winged morphs F-value = 244.67, p <2.2e-16).

Overall, these data indicate that there are considerable, long-term changes in gene expression among progeny in response to different embryonic environments, paralleling the observed impact of maternal density on morphology and reproduction (Figs. 1 and 2).

## Discussion

Our study shows that the maternal environment has long-lasting effects on embryos; effects that ultimately impact phenotypic integration in adulthood. We found that, at the morphological level, an adverse maternal environment decreases trait integration in progeny regardless of their morph. At the transcriptional level, we observed long-term signatures of the maternal environment in progeny of both morphs, although transcriptional integration decreased in response to an adverse environment in winged morphs but increased in wingless morphs. All of the morphological, reproductive, and transcriptional trait variation we observed was generated by a single genotype, demonstrating a remarkable ability of the environment to induce phenotypic variation even beyond the commonly considered winged versus wingless morph phenotypes.

Our study brings an explicit trait integration perspective to the field of intergenerational environmental effects. Many traits are affected by stressors across generations, and while the majority are parental effects (e.g., [7, 12, 58], some effects can persist for many generations [59–61]. This body of work has transformed our understanding of how environmental changes impact future generations, but most studies examined trait expression individually. The degree and pattern of trait integration can be context-dependent [62–64], and here this is what we observe. In pea aphids, the wingless and winged morphs are long-evolved responses to benign and stressful environments, respectively. We therefore expected each morph to be built of highly integrated units. And indeed, each morph displays evidence of morphological integration, especially as mediated by overall size (Fig. S3) but also with size effects removed (Fig. 2B, C). And both morphs show decreased patterns of integration in response to adverse maternal environments (Fig. 2A-C).

Our study also shows that different maternal treatments can result in large-scale changes in the adult transcriptional profiles of progeny, indicating a general re-patterning of physiology likely encoded by epigenetic changes initiated during embryogenesis. These effects are most apparent in the WGCNA analysis. The modules that differed in response to maternal treatment (Fig. 3D) together contained 4,601 genes (out of 9,845 assayed genes total), showing how impactful maternal environmental effects are in shaping adult physiology. GO enrichment of the significantly correlated gene sets revealed a range of different biological processes (Fig. 3D), again indicating that the long-term effects of maternal environment on progeny are systemic. One of the most gene-rich modules (the brown module with 1,556 genes) was enriched for metabolic processes, suggesting long-term effects of maternal environment on how the two morphs conduct energy production. Many of these transcriptional effects were also morph-specific, with more profound changes observed in the HD- versus LD-treated wingless morphs offspring compared to their winged sisters (Fig. 3A, 3D). This pattern is mirrored in the morphological trait response to maternal environment in wingless morphs, indicating that this morph responds more readily at multiple levels to environmental stimuli.

Given ongoing climate changes, there is increasing interest in predicting how species will respond to new patterns of environmental variation in terms of phenotypic plasticity ([43, 65–68]). To inform predictions, it is necessary to understand how traits co-vary at the level of the organism and at the level of the proximate mechanisms that generate covariation across traits. This is because how populations respond to existing or novel patterns of selection in heterogeneous environments depends on the degree to which traits exhibit plasticity independently or jointly, how new patterns of trait (co)variation align with the pattern of selection, and how developmentally flexible these relationships are [69–71]. Because we observed differences in trait integration in progeny across maternal environments, our study suggests that morphological traits may be more developmentally free to respond on a trait-by-trait basis under maternal stress as compared to benign conditions. The next logical steps would be to examine how genetic variation among pea aphids affects integration in alternative environments, and how longer, more intense, combined, or different types of stressors impact these patterns. Future work should explore whether the patterns we document extend to other systems, both with and without maternal stress-induced plasticity, as such plastic responses may confer greater developmental robustness.

## Acknowledgments

We thank Ryan Bickel and members of the Brisson lab for helpful discussion on this project. We thank Andy Jenson (AphidTrek) for specimen preservation and slide mounting protocol. This work was supported by the National Science Foundation under award number IOS 1749514 to JAB and Pumpprimer II funding from the University of Rochester.

## Notes

### Competing Interest Statement

The authors have declared no competing interest.

