## Supplementary figures and images for "Embryonic environment impacts adult traits and phenotypic integration in the pea aphid"

### Supplemental Figure 1

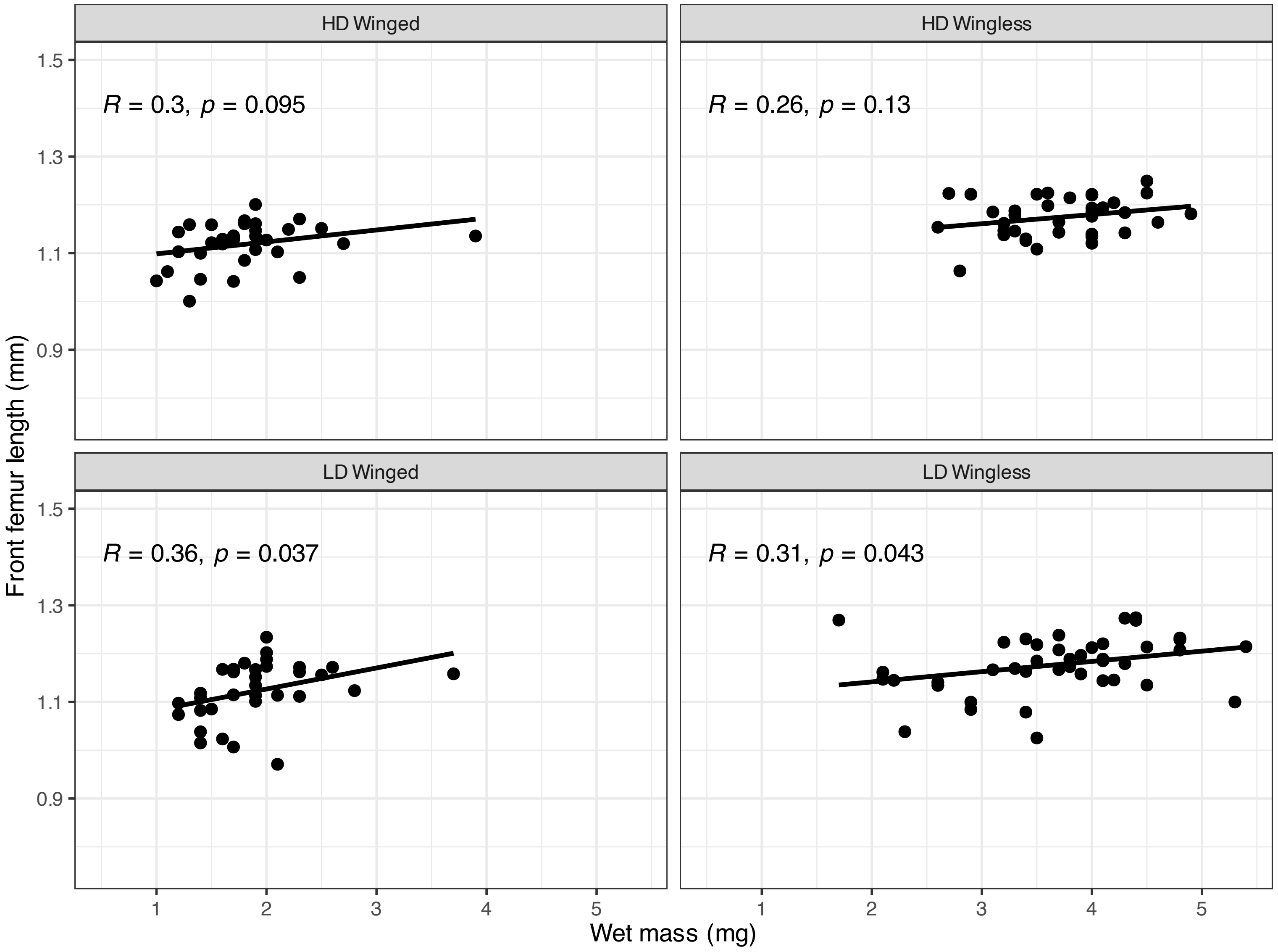

### Supplemental Figure 2

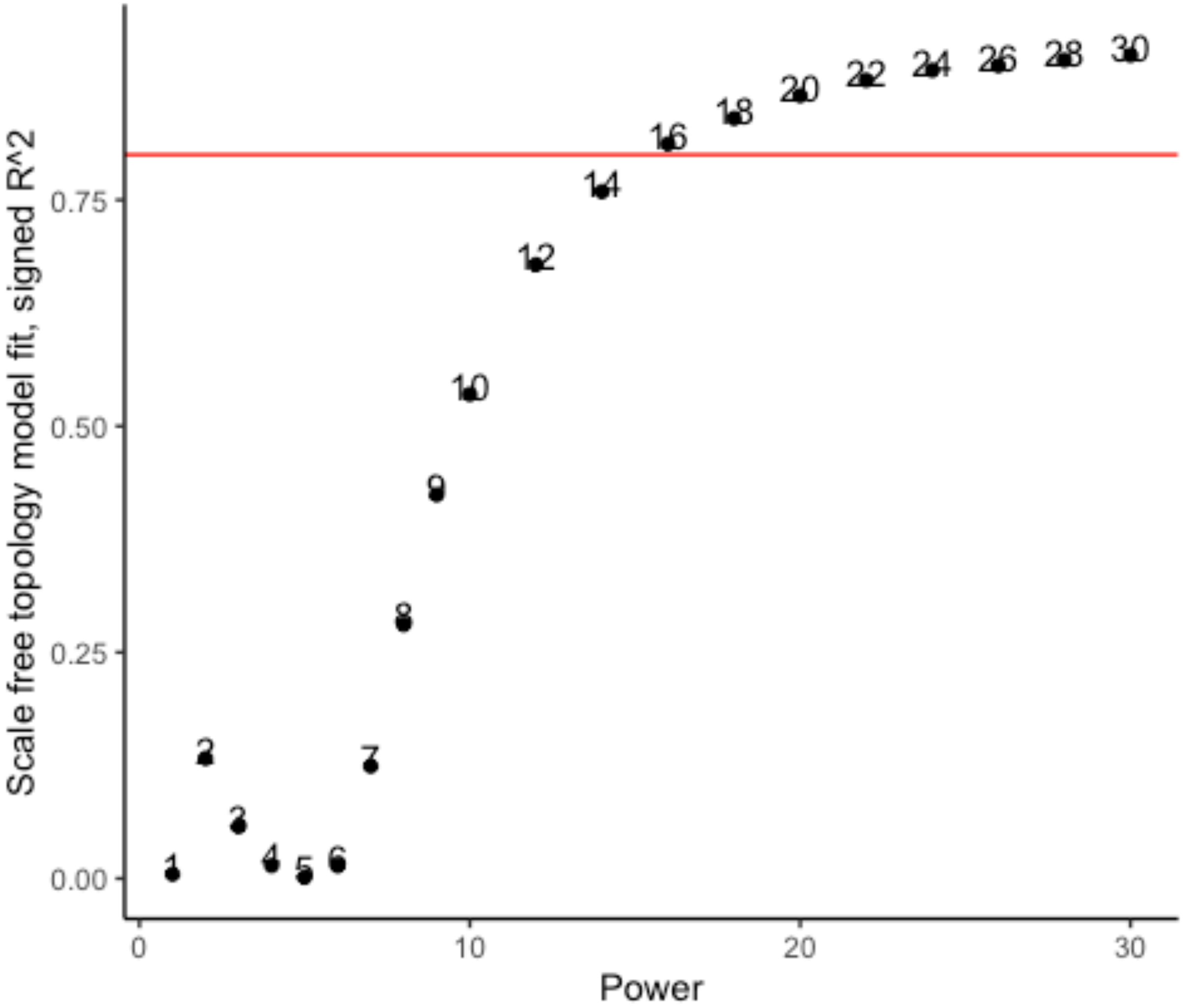

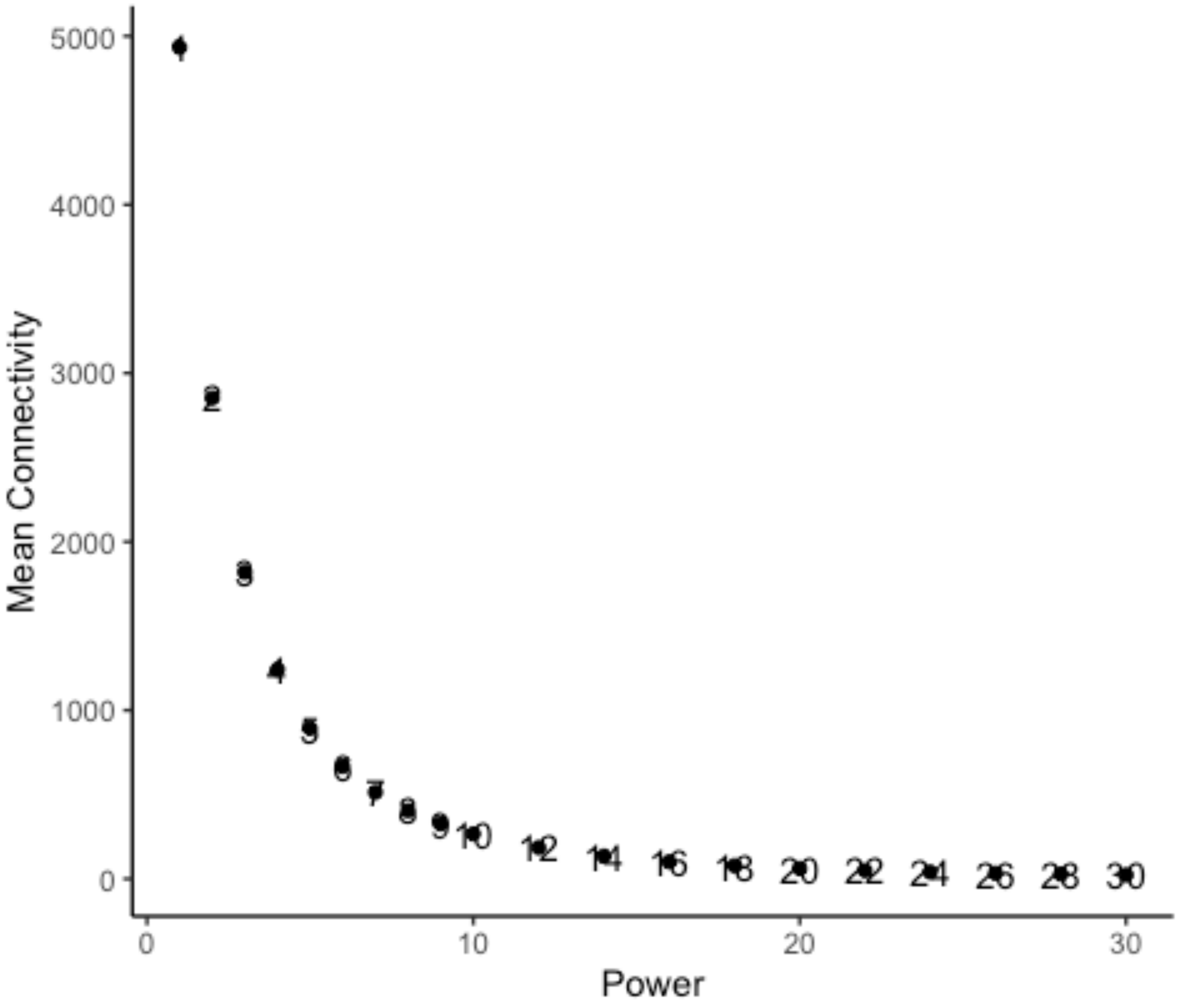

### Supplemental Figure 3

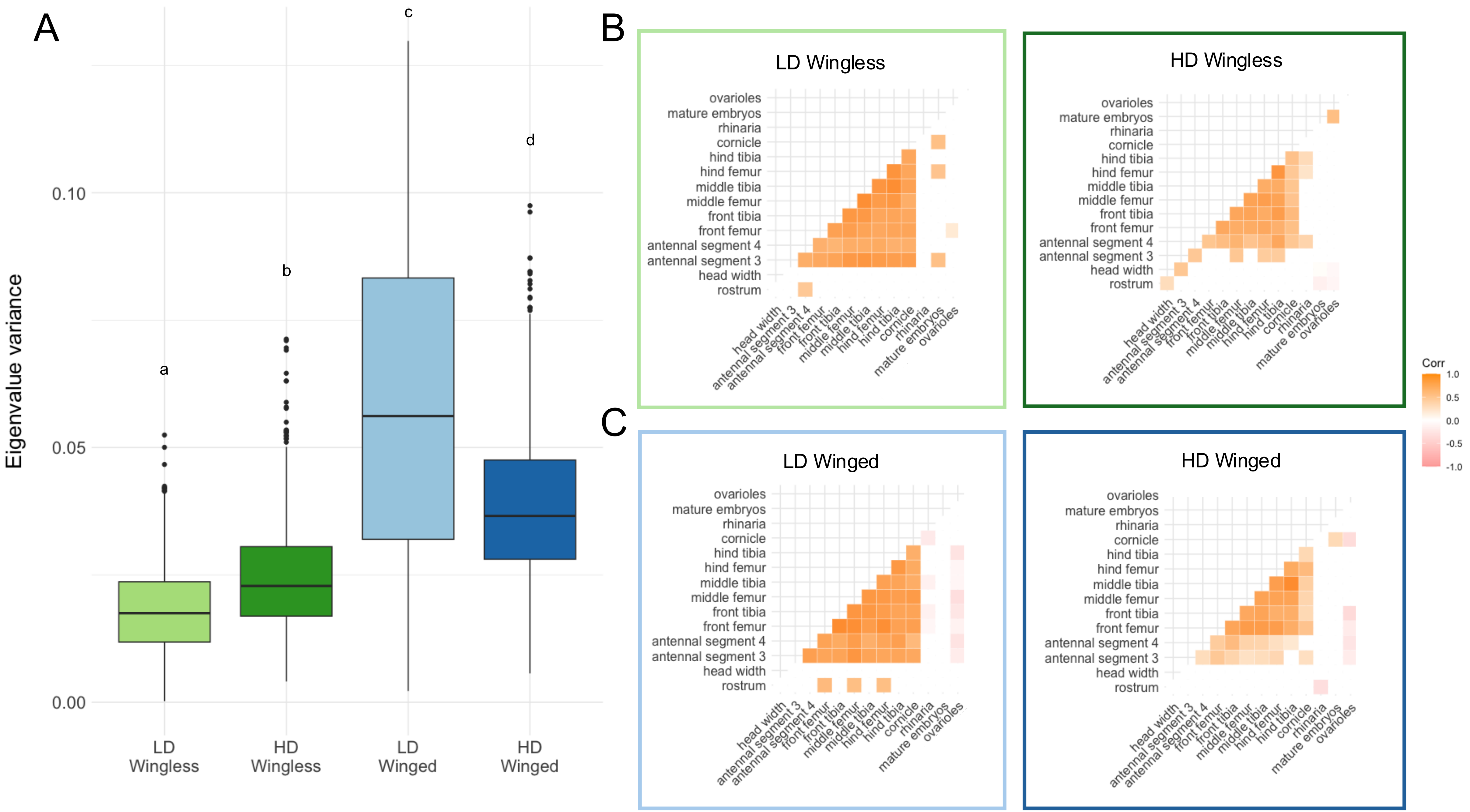

### Supplemental Figure 4

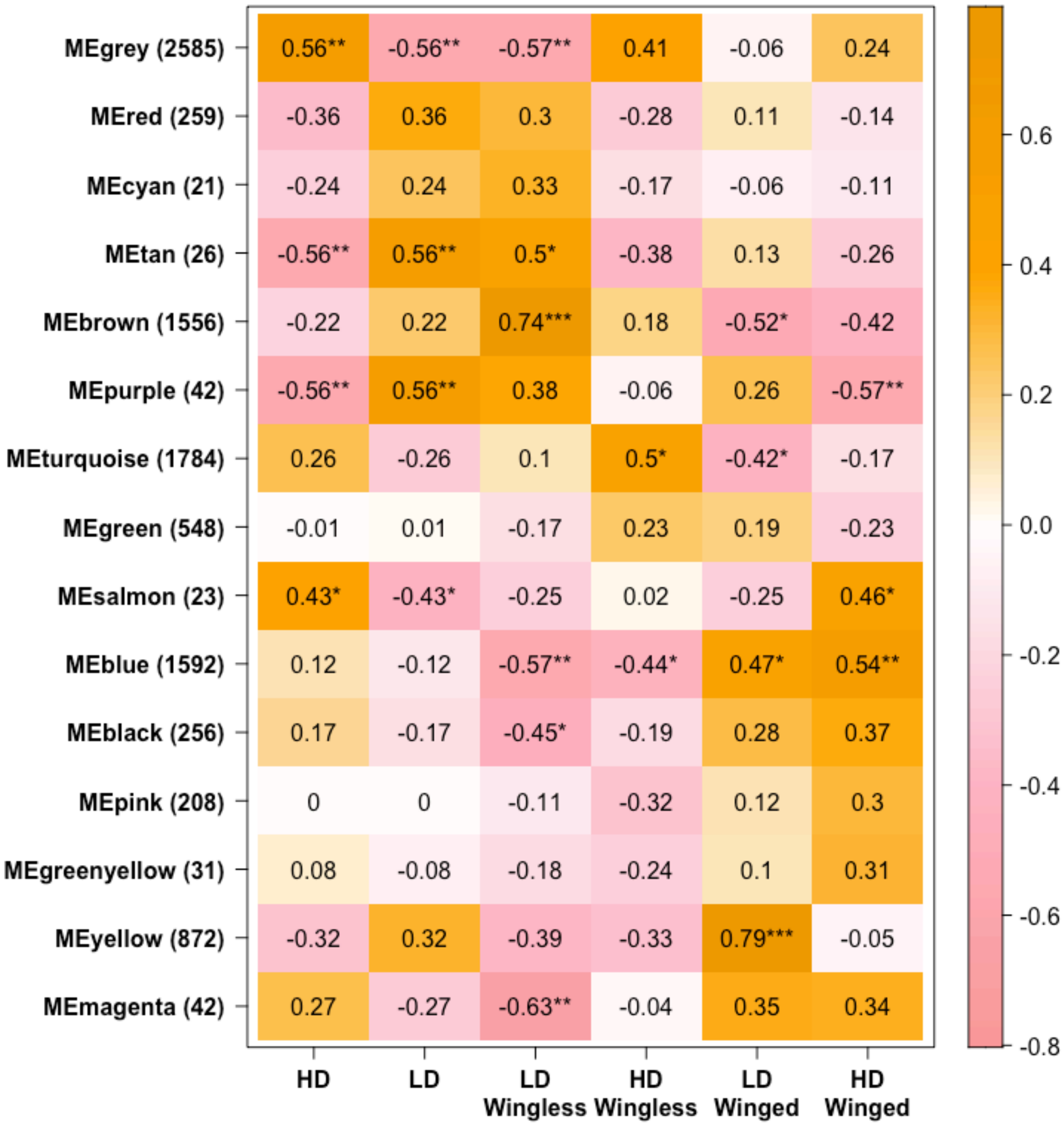
